# Reaction of Hydrogen sulfide homeostasis genes under biotic and abiotic stress condition in rice – computational approach

**DOI:** 10.1101/269639

**Authors:** Ganesh Alagarasan, Jegadeesan Ramalingam

## Abstract

Gaseous molecules are widespread signaling compounds, regulating the cell development process in all major plant parts. For many decades, hydrogen sulfide molecule is considered mainly for its deleterious effects on plant system. The increasing recent experimental evidence and phenomenal concepts on H_2_S molecule further advance our understanding of H_2_S interaction with plant tissues. In addition, the H_2_S messenger molecule is found to have positive effects on plant growth, in limited condition, to maintain the balanced homeostasis. To meet the increasing demand, and to sustain the crop yield, various crop improvement programs have been followed. However, there is a concern that traditional plant improvement method and increasing climate change has a negative impact on crop production. A major approach to combating plant stress is to evaluate and explore the alternate source mechanism(s). Towards this aim, it will be valuable to characterize the genes involved in H_2_S homeostasis in the staple food crop rice pan-genome. In this research, we identified 15 H_2_S homeostasis genes in rice and used it for the ~3k rice pan-genome analysis to find out the genetic relatedness based on single nucleotide polymorphism data. Multidimensional scale plot of 15 H_2_S homeostasis genes among the rice cultivars, and RNA-seq experimental data analysis under various biotic and abiotic stress shows the functional genes involved in biotic and abiotic stress. This study provides new insights into plant stress management in crop breeding and suggests how H_2_S gene(s) can be utilized to improve the agronomic traits in rice and other food crops.

## Introduction

Rice, maize, wheat and tapioca are important staple food crop across the world and these crops faces many challenges to attain its full genetic potential. Further, the combinatorial stress and standalone plant stress like salinity and drought are reducing the production and productivity potential of almost every food crop (Suzuki et al., 2014). Research projects with a focus on genetic and molecular analysis of unexplored functional genes will be very useful in crop breeding to combat the plant stress. Hence, the plant stress research has witnessed an increased attention, due to climate change, and continuous evolution of more virulent biotic and abiotic stress factor.

H_2_S homeostasis is an important source of mechanisms to tolerate the plant stress (Romero et al., 2014; Mostofa et al., 2015b; Dai et al., 2017; Tain et al., 2017). Recently, Mostofa et al. (2015b) reported the physiological implications of hydrogen sulfide in rice tissues under salinity stress. They have analyzed and discussed the effect of exogenous application of H_2_S and its effect on plant growth, particularly by maintaining a Na^+^/K^+^ ratio. The physiological functions of H_2_S in plants are mediated by sulfur-oxidation pathways (Mishanina et al., 2015), and different molecular targets, such as different ion channels, sulfate transporters and signaling proteins (Wang, 2012). In plants, the exogenous/endogenous hydrogen sulfide is responsible for conferring tolerance to both the biotic and abiotic stress (Bloem et al., 2004; Shi et al., 2015; Mostofa et al., 2015b). In every growth stage, plants do produce H_2_S in the cytosol through enzymatic mechanisms, particularly desulfhydrases. However, the H_2_S molecule also has the lethal effect on plant tissues at higher concentrations. Therefore, it is important to evaluate the potential of H_2_S gene in applied aspects before using it in crop breeding program. To explore the characteristic features and behavioral pattern of H_2_S homeostasis genes in biotic and abiotic stress, we have performed this study in rice H_2_S genes.

In this study, we focused to elucidate the role of H_2_S homeostasis genes under biotic and abiotic stress through combined computational genomics approach. Based on previous physiological experimental evidence (Mostofa et al., 2015a; Chen et al., 2017; Duan et al., 2015; Mostofa et al., 2015b), we have determined our gene identification and analysis criteria to charecterize the H_2_S homeostasis genes in rice. The reason for selecting the genes from a different category of hydrogen sulfide activity is mainly to maintain the balanced H_2_S content in plants. Since the higher H_2_S content will lead to tissue toxicity in rice, it is better to introgress, and/or clone the H_2_S homeostasis genes.

Despite the availability of multiple completely sequenced rice genomes, little is known on the occurrence of H_2_S in rice. Cultivated rice does not have all agriculturally desirable traits, which they might have lost during segmental, and/or tandem duplication events. As the wild rice contains many desirable traits, it is important to mine alleles from wild rice. ~3k rice genome project sequencing project made it possible to use the potential genes(s) from wide range of rice germplasm. Here we present the pan-genome analysis of the H_2_S homeostasis genes extended across the largest part of Oryza phylogeny using sequencing data from the 3k rice genome project.

## Materials and methods

### Comparative analysis of H_2_S homeostasis genes in rice

The keywords viz., sulfite reductase, cysteine synthase, cyanoalanine synthase and cysteine desulfyhdrase were searched in the Gramene and multiple rice database to retrieve the full-length gene sequences. Redundant results were filtered out for further downstream analyses. The retrieved gene sequences were manually annotated with FGENESH (http://www.softberry.com/). The annotated sequences were cross-checked in the public databases. The identified genes were positioned on their respective chromosome using the Oryzabase database. Number of intron and exons in the H_2_S genes were predicted in GSDS (http://gsds.cbi.pku.edu.cn/). The FGENESH derived protein sequences were subjected to conserved domain analysis in NCBI-CDD, Pfam identifiers, HMMER to confirm the presence H_2_S homeostasis domains. These protein sequences were used for Phylogenetic classification of H_2_S homeostasis genes through W-IO-TREE (http://iqtree.cibiv.univie.ac.at/) server with default parameters. Protein topology and signal peptides were predicted through Protter. To find the potential miRNA targets, the gene sequences were scanned in psRNATarget (http://plantgrn.noble.org/psRNATarget/) server.

The association between the potential miRNA expression with agronomic traits was done in RiceATM (http://syslab3.nchu.edu.tw/rice/). Rice pan-genome analysis was performed in the rice SNP seek database (http://snp-seek.irri.org/) to determine the genetic distance of ~3k rice genotypes based on H_2_S genes. For promoter analysis, 2 kb upstream of all gene sequences was subjected to transcription factor analysis in PlantPAN database. Whole genome RNA-seq data were used to determine the expression of fifteen H_2_S in various abiotic stress phosphorus stress (PRJEB11899); drought stress and salinity stress (GSE60287). To compare the H_2_S gene expression level in biotic stress, bacterial blight (GSE57670) transcriptome data were used to generate the FPKM value. The obtained FPKM value was used to generate the heat map to quantify the transcript abundance.

## Results

### *Insilico* functional characterization of H_2_S gene family

To comprehensively investigate and characterize the H_2_S gene family in rice, a genome-wide survey covering the entire length of all the 12 rice chromosomes were performed. The hidden Markov model and keyword-based and search in Gramene and rice genome annotation database resulted in the identification of 15 potential full-length H_2_S homeostasis related genes. The H_2_S homeostasis genes identified in the rice genome database and their features are mentioned in (Table). The chromosomal localization analysis of rice H_2_S genes revealed variable distribution of the genes in all chromosomes except for chromosome 7, 8, 9, 10 and 11. While a maximum number of four genes were located on chromosome 1 and 6. In contrast, only one gene was identified in both the chromosome number 4 and 12. Of fifteen H_2_S genes, three (OsH_2_S12, OsH_2_S13 and OsH_2_S14) were defined as sulfite reductase, ten (OsH_2_S2, OsH_2_S3, OsH_2_S4, OsH_2_S5, OsH_2_S6, OsH_2_S7, OsH_2_S8, OsH_2_S9, OsH_2_S10 and OsH_2_S15) were defined as cysteine synthase and one as (OsH_2_S1) cyanoalanine synthase and another one as (OsH_2_S11) L-cysteine desulfydrase. The H_2_S homeostasis family genes posses a small number of introns in their sequences (Figure 2).

Based on the results of Pfam, HMMER, CDD and Phylogenetic classification of conserved domain analysis, the H_2_S homeostasis genes were independently grouped as sulfite reductase, cysteine synthase, cyanoalanine synthase and cysteine desulfyhdrase (Figure 1). Bzip, NAC, WRKY, MYB and AP2/EREBP are the predominant stress related transcription factor present in all the H_2_S genes promoter sequence. Of 15 H_2_S genes, only two genes (OsH_2_S2 and OsH_2_S14) had a potential miRNA binding site (Figure 1). In total, seven miRNA (osa-miR818a, osa-miR818b, osa-miR818c, osa-miR818d, osa-miR818e, osa-miR1436 and osa-miR2879) targeting two genes have been identified. The miRNA (osa-miR818a, osa-miR818b, osa-miR818c and osa-miR818e) are found to strongly associated with the plant height and 1000 seed grain weight (osa-miR818b and osa-miR818b). In the surveyed ~3k rice accessions, 2757 accessions have all the 15 H_2_S homeostasis genes. The genes OsH_2_S3, OsH_2_S12 and OsH_2_S15 had a number of allelic variations across the surveyed rice pangenome. The pan-genome analysis revealed a good genetic diversity analysis based on H_2_S sequences. This may help in selecting the donor plants with potential H_2_S alleles in plant breeding programs. More number of indica type rice possess all the 15 genes, while the japonica rice has the second most number H_2_S genes.

**Figure 1:**
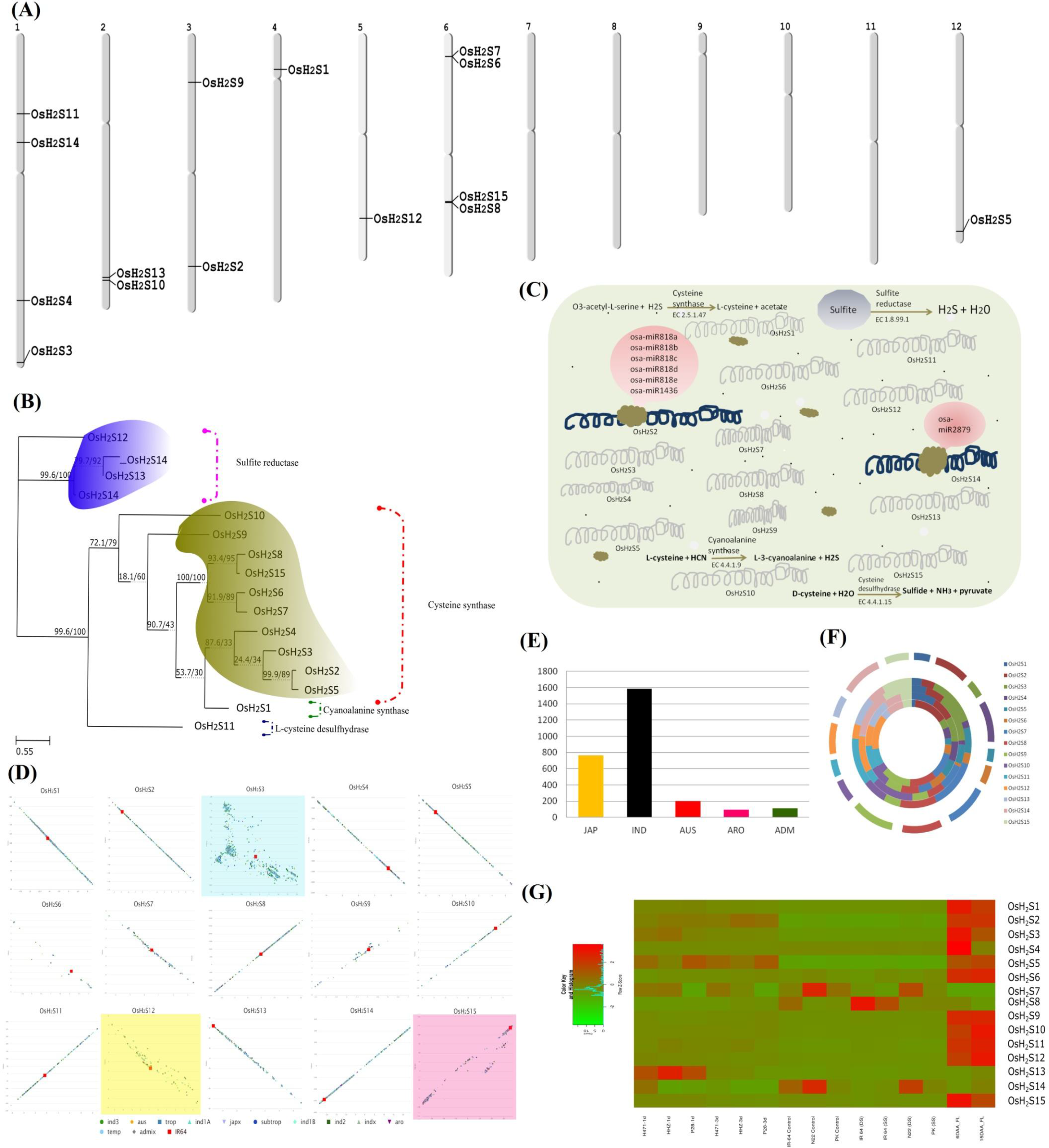
Genome-wide characterization of H_2_S homeostasis genes in rice. **(A)** Chromosomal positioning and distribution of genes. H_2_S genes were identified using a combined computational approaches in rice, i.e., key word search, conserved domain identification, Pfam identifier and HMMER search. Chromosomal positioning was based on the physical position (Mb) in 12 rice chromosomes. The chromosome number is indicated at the top of each chromosome. **(B)** The evolutionary history was inferred by using the Maximum Likelihood method based on W-IQ-TREE. The bootstrap consensus tree inferred from 1000 replicates is taken to represent the evolutionary history of the taxa analyzed (Felsenstein, 1985). The resulting four major clusters were shown in the figure. The analysis involved 15 amino acid sequences. All amino acid sequences in this study have been manually annotated in FGENESH to avoid the redundancy. **(C)** miRNA scanning and target prediction. The full-length nucleotide sequences were subjected to verify the presence of miRNA targets, Of 15 H_2_S gene, only two gene sequence had a potential miRNA binding site. Interestingly, the gene OsH_2_S2 has a single binding site for six marinas. **(D)** Pan-genome analysis in ~3k rice genome sequence data. The pan-genome analysis revealed the possible and potential H_2_S alleles across wide rice accessions. The data in this study are obtained from Rice SNP seek database. The genetic relatedness among these accessions is drawn based on the variations in any particular H_2_S gene. IR-64 variety is highlighted in gene based genetic diversity analysis **(E)** The distribution of all 15 H_2_S homeostasis genes in wide rice accessions. The graph indicates the number of any particular rice type having all 15 H_2_S. The results show that maximum number of *Indica* type rice possess all H_2_S homeostasis genes. **(F)** The distribution of stress related transcription factors in the 2kb upstream nucleotide sequences. The five rings in the figure indicate the five transcription factor. From outer side, ring 1− Bzip, ring 2− NAC, ring 3− WRKY, ring 4− MYB and ring 5− AP2/EREBP. **(G)** Functional characterization of H_2_S homeostasis genes. Heatmap showing the expression of H_2_S genes under salinity and drought stress. The FPKM value is calculated from the RNA-seq data derived from the whole genome transcriptome study under salinity and drought stress. The colored bar at the left indicates the relative expression value, wherein, −2.0, 0.0 and 2.0 indicates low, medium and high expression respectively.

To check the expression level of all 15 genes under P stress, salinity & drought stress and bacterial blight stress, the RNA-seq were analyzed (Figure 2). The differential expression pattern of H_2_S genes strongly supports the potential role of H_2_S homeostasis genes under various biotic and abiotic stress. For example, in IR 64 variety the cysteine synthase genes (OsH_2_S8 and OsH_2_S9) were up-regulated under drought stress. In Pokkali OsH_2_S1, OsH_2_S6, OsH_2_S10, OsH_2_S13 were up-regulated and OsH_2_S4, OsH_2_S5, OsH_2_S12 and OsH_2_S15 were highly down-regulated under salinity stress. Under biotic stress condition (bacterial blight), the genes OsH_2_S1, OsH_2_S3, OsH_2_S5 and OsH_2_S12 had a maximum level of expression. While in P stress, the genes OsH_2_S1, OsH_2_S3, OsH_2_S5, OsH_2_S12 and OsH_2_S15 had a maximum expression and OsH_2_S7 had a negligible/or low expression level. Among biotic and abiotic stress condition, the H_2_S genes expression is comparatively higher in P stress condition. Hence the research should be focused more on characterizing the H_2_S homeostasis genes under various P treatments in rice.

**Figure 2:**
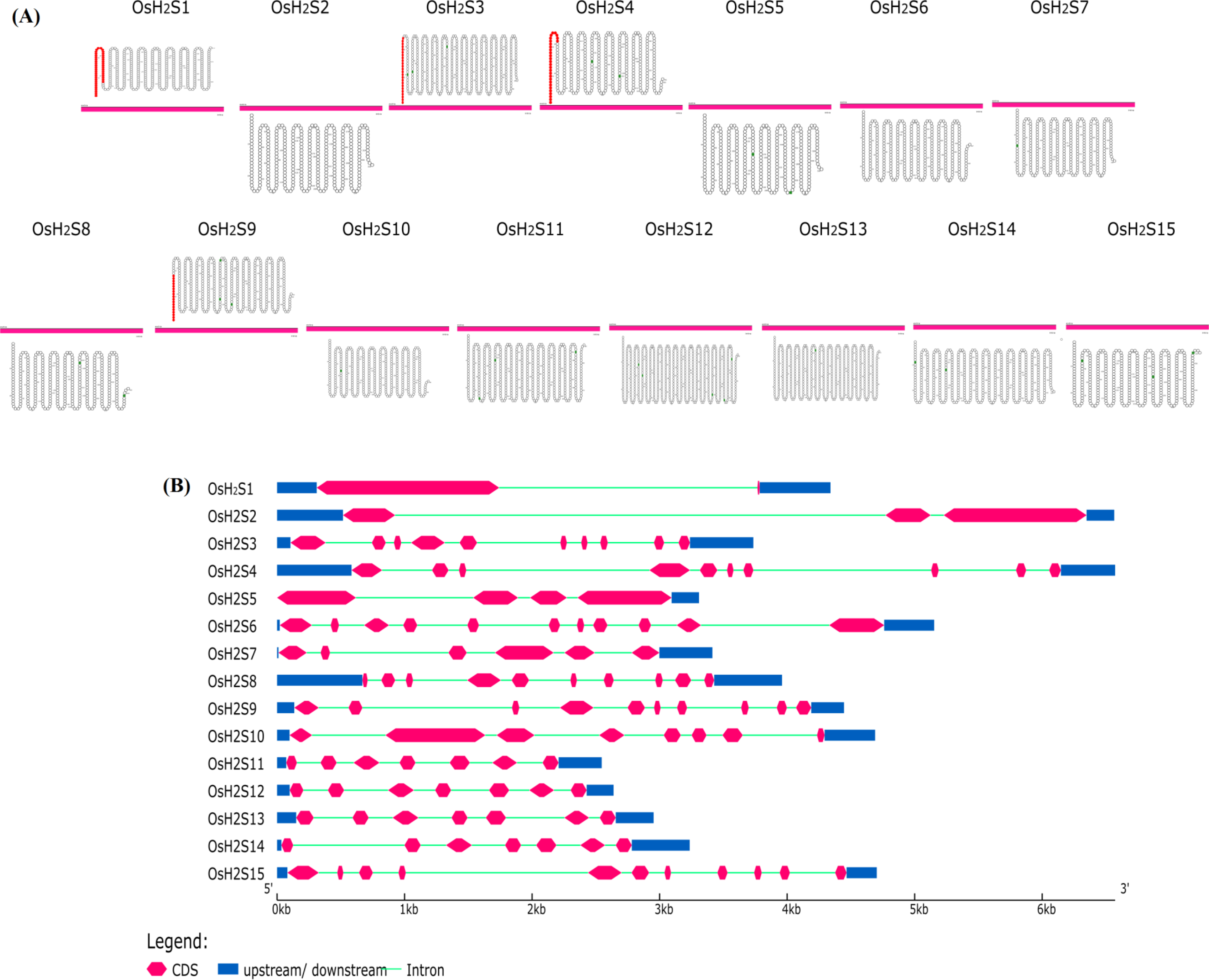
H_2_S protein membrane topology and gene structure. **(A)** Signal peptide prediction and orientations of membrane-spanning segments with respect to the inner and outer sides of the plant cell plasma membrane. The pink colored peptide chain at the end of N-terminus indicates the presence of signal-peptide in the H_2_S homeostasis gene **(B)** representation of the presence and arrangements of number of introns/exons in the genes.

## Conclusion

In this paper, an in-depth *insilico* gene characterization of the H_2_S family of rice was performed. We identified and characterized 15 H_2_S homeostasis genes in rice. The phylogenetic grouping of protein sequences confirmed the presence of conserved domains in the H_2_S related gene family. In addition, the presence of H_2_S gene family in seven chromosomes reflects the unequal distribution in the rice genome. Analysis of the promoter sequence, transcription factors and quantification of transcript abundance enabled the retrieval of valuable information related to the functional response, diversity in the stress responsive elements and biotic/abiotic stress responsiveness of these genes. Finally, a comparative analysis between rice accessions revealed a high degree of sequence conservation/variation within the H_2_S domain as well as in the domain organization of these genes. Furthermore, analysis of the expression profiles of the H_2_S genes confirmed that they are differentially regulated in response to several types of stress. These data suggest a potential role for the H_2_Ss in plant signaling and defense mechanisms.

### Future direction of research

Interestingly, Neale et al., (2017) reported that H_2_S signals produced from the plants have the ability to alter pathogenecity of microbes. The interaction between H_2_S genes and microbes, whether the triggered plant H_2_S genes are race-specific or race non-specific, and the genes and/or QTL controlling the specificity are needs to be clearly addressed. Research describing the reaction of plant H_2_S genes with specific microbial receptor protein will reflect the outcome of the interactions between alleles at all avirulence loci in the phyto-pathogen and alleles at all H_2_S loci of the plant gene. In addition, delineating the role of H_2_S directed regulation of abiotic stress responsive genes/QTLs/transcription factors will provide clues to the mechanisms controlling H_2_S homeostasis in plants. Further, the application of next-generation sequencing techniques will explore the presence of genotype specific novel INDEL region/SNPs in the H_2_S genes in plants. Some of the H_2_S genes have the signal peptide and are predicted to be involved in the secretory pathway. It would be interesting to see whether these signal peptides have any role in protein targeting and what happens if we truncate the signal peptide. It will also be motivating to observe the localization pattern of H_2_S proteins. The current evidences on the role of signal peptides suggests that these proteins are secreted in some other cellular components, and being transported to inter-cellular spaces. Meanwhile, H_2_S genes have the potential to interact with other stress related genes (Wang, 2012). In addition, hydrogen sulfide has a positive effect on plant growth at various stress condition (Chen et al., 2013; Christou et al., 2013; Li et al., 2012; Suzuki et al., 2014; Wang et al., 2010; Wang et al., 2012; Zhang et al., 2010a; Zhang et al., 2010b; Zhang et al., 2010c). Hence, the similarities of these longer genes should be better studied in order to analyze their significance in altering plant tolerance to stress and/or other important agronomic traits that may bring interesting insights about H_2_S evolution or that may be of interest of plant breeders.

### Author Contributions

GA and JR initiated the project. All the authors have made a substantial, direct, intellectual contribution to the work, and reviewed the final version of the manuscript.

## Acknowledgments

The authors acknowledge the assistance from Dr. Abdul Baten from Southern Cross Plant Science, Southern Cross University, Lismore, NSW, Australia in analyzing RNA-seq derived expression patterns.

